# Cell-type-dependent bioelectric regulation of fin size and fin-ray segment length in zebrafish

**DOI:** 10.1101/2024.03.27.587111

**Authors:** Toshihiro Aramaki, Shigeru Kondo

## Abstract

During development, organ size and the size of its components must be coordinated. However, the mechanisms integrating these two growth processes remain poorly understood. Here, using the zebrafish fin as a model, we investigated how bioelectric signaling regulates overall appendage size and fin-ray bone segment length. To manipulate bioelectric signaling, we expressed the hyperactive potassium channel mutant kcnk5b^W169L^ or the dominant-negative connexin mutant Cx43^T154A^ under a common set of cell-type-specific promoters. Cell-type-specific bioelectric manipulation in keratinocytes primarily affected fin size, whereas manipulation in osteoblasts preferentially altered fin-ray segment length. Combining these manipulations generated enlarged fins with short segments or small fins with long segments, providing direct evidence that fin size and fin-ray segment length can be regulated independently. Furthermore, optogenetic manipulation demonstrated that membrane potential itself can regulate fin size and fin-ray segment length. Together, these findings reveal that bioelectric signaling can independently regulate appendage growth and skeletal element growth through distinct cellular targets.

**SUMMARY STATEMENT:** Bioelectric signaling in distinct cell types independently influences appendage size and bone segment length, revealing separable regulation of organ growth and skeletal patterning.

## INTRODUCTION

How organ growth is coordinated with that of its component tissues remains a fundamental question in developmental biology. In vertebrates, overall appendage size is closely associated with the size of skeletal elements that provide structural support, yet how these two growth processes are coordinated remains incompletely understood. Zebrafish fins have emerged as a powerful model for investigating appendage growth. During postembryonic growth, fin size is scaled to body size (Iovine and Johnson, 2000), while regenerating fins faithfully restore their original size following amputation (Kujawski et al., 2014; Pfefferli and Jaźwińska, 2015). Teleost fins are supported by many rod-shaped skeletal structures called fin rays (Grandel and Schulte-Merker, 1998; Pfefferli and Jaźwińska, 2015). Each fin-ray consists of tandemly repeated bony segments whose lengths are also tightly controlled (Iovine and Johnson, 2000). Together, these characteristics provide an opportunity to investigate how organ-level fin growth is coordinated with the growth of its component skeletal elements.

Genetic studies have established bioelectric signaling as a major regulator of fin growth. In the *another-long-fin* (*alf*) mutant, a gain-of-function mutation in the potassium channel kcnk5b produces enlarged fins with elongated fin-ray segments (Haffter et al., 1996; Perathoner et al., 2014; van Eeden et al., 1996) (Fig. 1A, B; Fig. S1). By contrast, in the *short-fin* (*sof*) mutant, loss of gap junction protein connexin43 (cx43) function reduces both overall fin size and segment length (Iovine and Johnson, 2000; Iovine et al., 2005) (Fig. 1C). Interestingly, the *longfin* (*lof*) mutant, caused by ectopic expression of the potassium channel kcnh2a, exhibits fin overgrowth without elongation of fin-ray segments (Iovine and Johnson, 2000; Stewart et al., 2021). Together, these findings suggest that different bioelectric regulators (ion channels) may uncouple organ growth from skeletal growth, raising the fundamental question of how these distinct fin growth patterns are specified.

**Figure 1.**
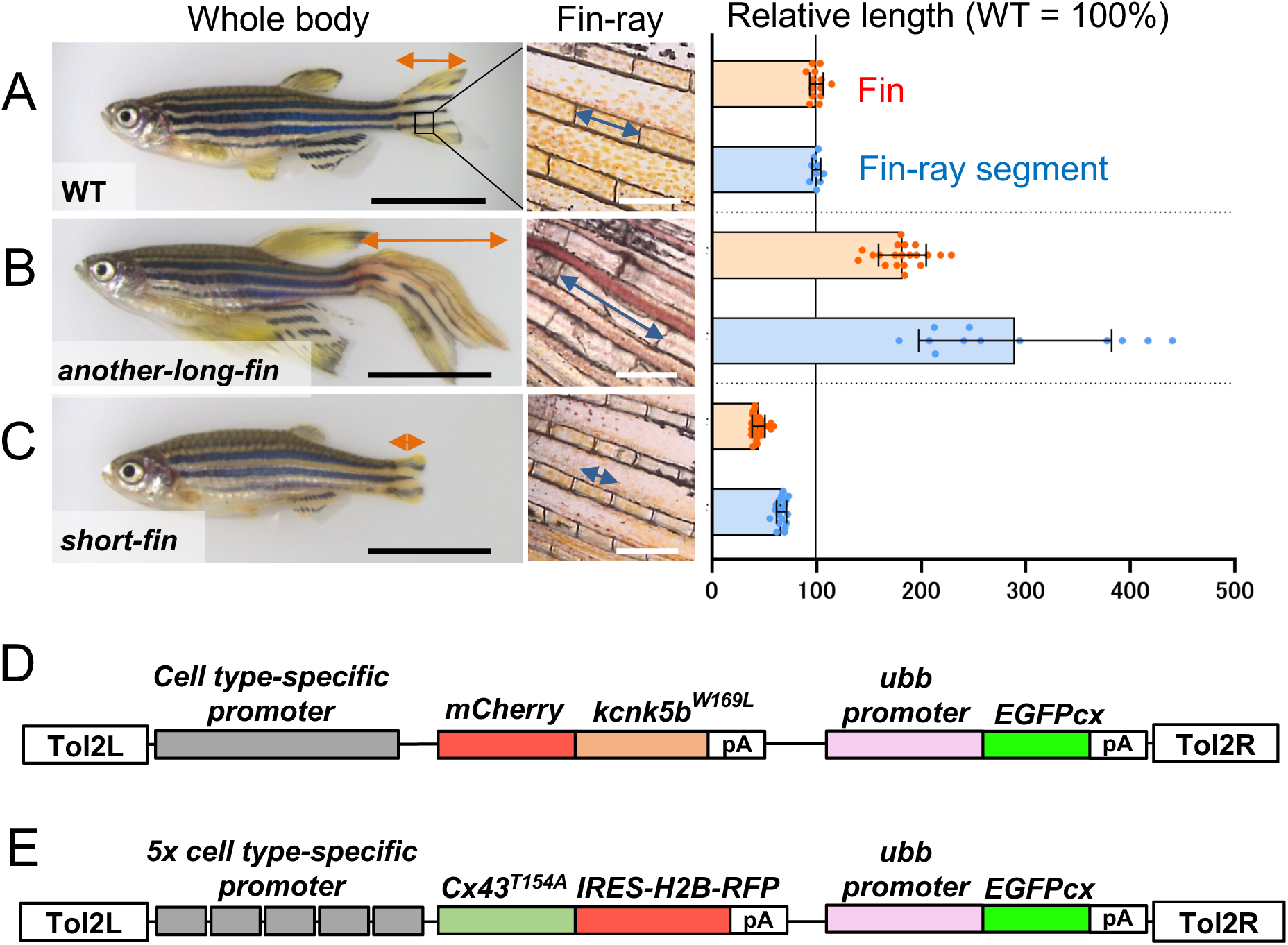
A transgenic screening strategy for identifying cell types regulating fin morphology. (A-C) Zebrafish mutants with abnormal fin morphology. Wild-type fish (A), the *another-long-fin* mutant with expanded fins and elongated fin-ray bone segments (B), and the *short-fin* mutant with reduced fins and shortened bone segments (C). Black scale bars in whole body images, 10 mm. White scale bars in fin-ray images, 300 µm. Plots on the right show caudal fin length and fin-ray segment length relative to wild type. Data are presented as mean ± SD (n = 9–26 fish per group). Mean values for wild-type fish were set to 100. (D, E) Schematic diagrams of transgenic constructs used for promoter screening.

To understand how these different growth patterns are generated, it is essential to identify the cellular targets of each bioelectric regulator. Although previous studies have implicated closely related mesenchymal cell populations in the actions of kcnk5b, cx43, and kcnh2a, these conclusions were reached using different experimental approaches, including cell transplantation, inducible transgene expression, and expression analyses (Dardis et al., 2017; Perathoner et al., 2014; Stewart et al., 2021). Because each regulator was investigated independently, it has remained unclear whether these regulators act through a common mesenchymal cell population or through distinct cellular targets.

Here, to address this question, we artificially reproduced mutant phenotypes by expressing a hyperactive kcnk5b mutant or a dominant-negative cx43 mutant under the control of a common set of cell-type-specific promoters. Using a unified experimental framework, we directly compared the cellular targets of two representative bioelectric regulators, allowing us to determine whether fin size and fin-ray segment length are controlled through common or distinct cellular populations.

## RESULTS

### Cell-type-specific transgenic screening identifies cellular targets of bioelectric regulators

To identify the cellular targets of bioelectric regulators, we performed a cell-type-specific transgenic screen using the Tol2 transposon system (Fig. 1D, E). We expressed a hyperactive kcnk5b mutant (kcnk5b^W169L^) (Perathoner et al., 2014) or a dominant-negative rat Cx43 mutant (Cx43^T154A^) (Beahm et al., 2006) under the control of multiple cell-type-specific promoters (Table 1; Fig. S2, S3). For effective inhibition by a dominant-negative mutant, we used five tandem copies of each promoter (5x promoter) to enhance transgene expression (Fig. S4, S5). This was considered particularly important for Cx43^T154A^ because connexins assemble into hexameric channels (connexons) (Beahm et al., 2006; Sáez et al., 2003). Among the promoters examined, transgenes driven by the *keratin4* (*krt4*), *RUNX2* and *osterix* (*osx*) promoters reproducibly recapitulated the phenotypes of *another-long-fin* and *short-fin* mutants. In contrast, *actinodin1^MC^* promoter-driven transgenes promoted fin overgrowth but did not reproduce the corresponding short-fin phenotype (Table 1).

**Table 1.** Cell type screening using tissue-specific promoters. In the first screening with kcnk5b^W169L^ transgenes, the presence (+) or absence (−) of *another-long-fin*-like phenotypes was evaluated in F0 founder fish. In the second screening, Cx43^T154A^ transgenes were tested only using promoters that reproduced the *another-long-fin* phenotype in the first screening. The presence (+) or absence (−) of *short-fin*-like phenotypes was evaluated in F1 stable transgenic fish. NT, not tested.

| Promoter | Cell type | kcnk5b<sup>W169L</sup> Tg (F0) |  | Cx43<sup>T154A</sup> Tg (F1) |  |
| --- | --- | --- | --- | --- | --- |
|  |  | Larger fin | Longer bone | Smaller fin | Shorter bone |
| *ubiquitin-b* | ubiquitous | + | + | NT | NT |
| *keratin1c19e* | basal-keratinocyte | - | - | NT | NT |
| *keratin4* | outer-keratinocyte | + | - | + | - |
| *collagen2a1a* | chondrocyte | - | - | NT | NT |
| *RUNX2* | osteoblast early | - | + | - | + |
| *osterix* | osteoblast middle | - | + | - | + |
| *osteocalcin* | osteoblast late | - | - | NT | NT |
| *actinodin1<sup>MC</sup>* | mesenchymal cell | + | - | - | - |
| *msxb* | mesenchymal cell | - | - | NT | NT |
| *acp5a* | osteoclast | - | - | NT | NT |

The screening identified distinct cellular targets underlying fin size and fin-ray segment length regulation. Keratinocyte-specific *krt4* promoter-driven transgenes preferentially reproduced fin size phenotypes, whereas osteoblast-specific *RUNX2* and *osx* promoter-driven transgenes preferentially reproduced fin-ray segment phenotypes (Table 1). These results suggest that fin size and fin-ray segment length may be regulated independently through distinct cell populations. Therefore, we focused subsequent analyses on these keratinocyte- and osteoblast-specific transgenic lines.

### Keratinocytes regulate fin size, whereas osteoblasts regulate fin-ray segment length

To confirm the cellular targets identified in the screening, we established stable transgenic lines and analyzed their phenotypes in detail. These transgenic fish showed cell-type-specific expression patterns driven by the corresponding promoters (Fig. S3, S5). Keratinocyte-specific expression of kcnk5b^W169L^ markedly expanded fins with only a slight effect on fin-ray segment length (Fig. 2A), whereas keratinocyte-specific expression of Cx43^T154A^ reduced fin size without affecting segment length (Fig. 2B). These fin size changes were observed in all fins (Fig. S6). In contrast, osteoblast-specific expression of kcnk5b^W169L^ preferentially elongated fin-ray segments (Fig. 2C), whereas osteoblast-specific expression of Cx43^T154A^ selectively shortened them with only a modest effect on overall fin size (Fig. 2D). Collectively, these results demonstrate that the same bioelectric regulators produce distinct developmental outcomes depending on their cellular targets.

**Figure 2.**
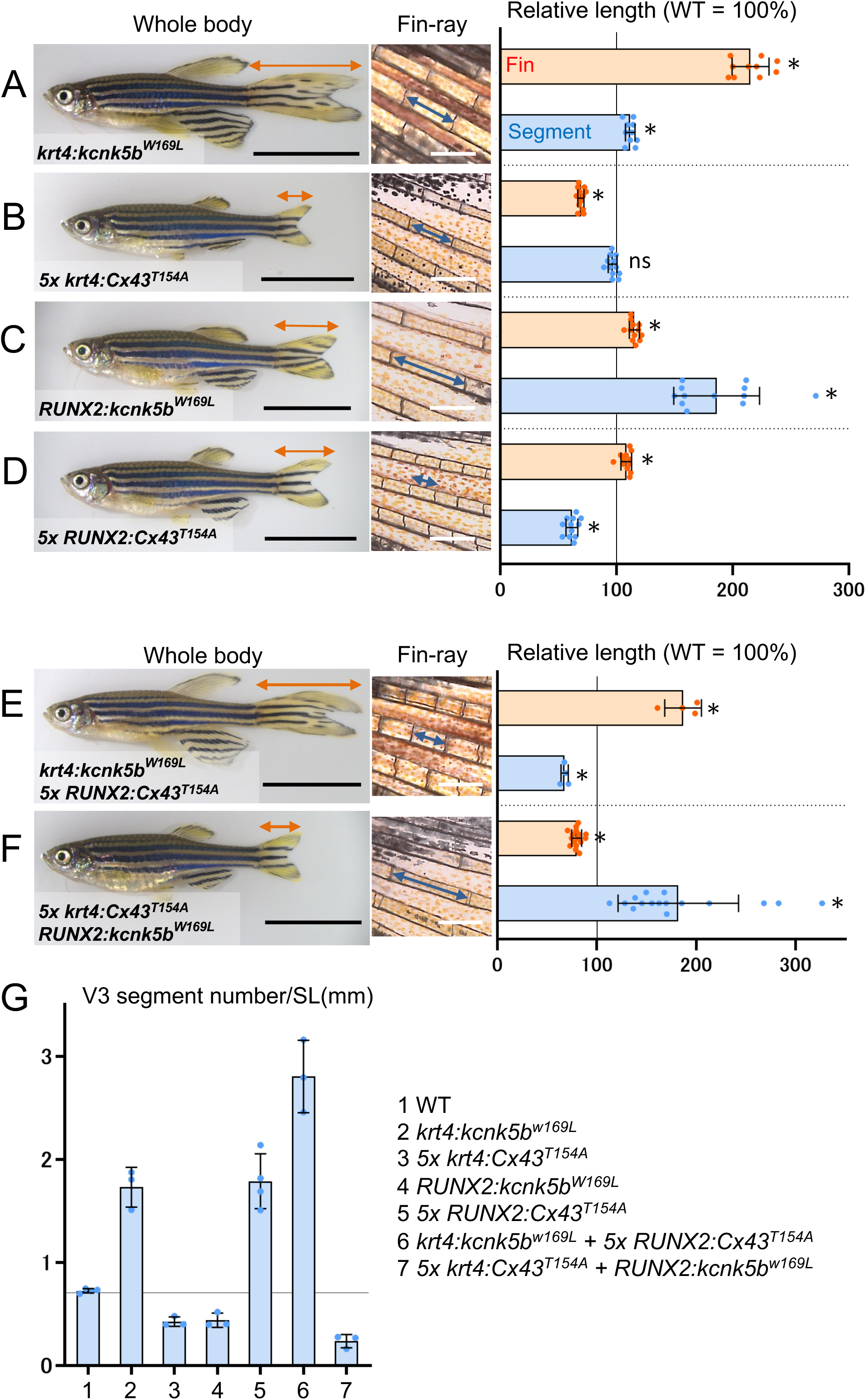
Independent regulation of fin size and fin-ray segment length by cell-type-specific bioelectric manipulation. (A-F) Stable transgenic fish recapitulating fin size or fin-ray segment length phenotypes. (A) *krt4:kcnk5b^W169L^* transgenic fish expressing kcnk5b^W169L^ in keratinocytes. (B) *5x krt4:Cx43^T154A^* transgenic fish expressing Cx43^T154A^ in keratinocytes. (C) *RUNX2:kcnk5b^W169L^* transgenic fish expressing kcnk5b^W169L^ in osteoblasts. (D) *5x RUNX2:Cx43^T154A^* transgenic fish expressing Cx43^T154A^ in osteoblasts. (E) Double-transgenic fish expressing kcnk5b^W169L^ in keratinocytes and Cx43^T154A^ in osteoblasts. (F) Double-transgenic fish expressing Cx43^T154A^ in keratinocytes and kcnk5b^W169L^ in osteoblasts. Data are presented as mean ± SD (n = 4–16 fish per group). The mean values for WT fish were set to 100. *P < 0.05 versus WT (Welch’s one-way ANOVA followed by Dunnett’s multiple-comparisons test). ns, not significant. Black scale bars in whole-body images, 10 mm. White scale bars in fin-ray images, 300 µm. (G) Number of segments in the ventral third (V3) fin ray normalized to standard length (Fig. S1). Data are presented as mean ± SD (n = 3–4 fish per group).

We next asked whether fin size and fin-ray segment length could be manipulated independently within the same individual. Double-transgenic fish expressing kcnk5b^W169L^ in keratinocytes together with Cx43^T154A^ in osteoblasts developed enlarged fins composed of short fin-ray segments, resulting in an increased number of segments (Fig. 2E, G lane 6). In contrast, the reciprocal combination generated small fins with longer fin-ray segments, resulting in a reduced number of segments (Fig. 2F, G lane 7). These findings provide direct evidence that fin size and fin-ray segment length can be regulated independently through distinct cellular targets, despite involving the same bioelectric regulators.

### Cell type-specific rescue reveals distinct requirements for fin size and fin-ray segment length regulation

To further examine the functional relevance of the cellular targets identified in the screening, we tested whether cell-type-specific bioelectric manipulation could rescue the mutant phenotypes. In the *alf* mutant, expression of dominant-negative kcnk5b (Perathoner et al., 2014) in keratinocytes did not rescue either the fin size or the fin-ray segment length (Fig. 3A, B). This result suggests that bioelectric manipulation of keratinocytes alone is insufficient to rescue fin overgrowth in the *alf* mutant, implying that additional cell populations contribute to fin size regulation. In contrast, expression of dominant-negative kcnk5b in osteoblasts selectively rescued the elongated fin-ray segment phenotype without affecting fin size (Fig. 3C), indicating that bioelectric manipulation in osteoblasts is sufficient to rescue the fin-ray segment elongation in the *alf* mutant.

**Figure 3.**
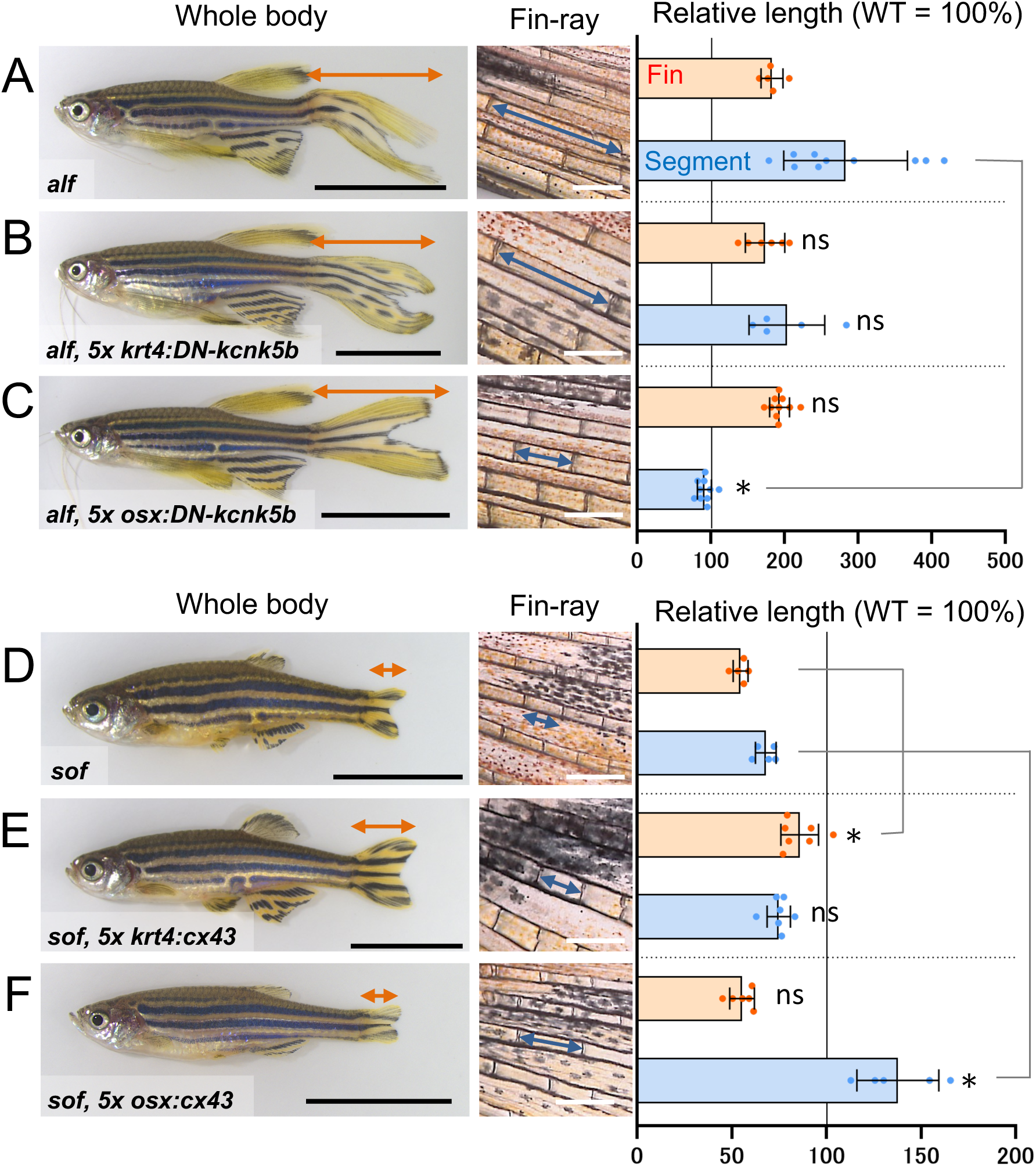
Cell-type-specific rescue of fin mutant phenotypes. (A-C) Rescue experiments for the *another-long-fin* mutant phenotype. (A) *alf* mutant. (B) *alf* mutant expressing dominant-negative kcnk5b in keratinocytes. (C) *alf* mutant expressing dominant-negative kcnk5b in osteoblasts. (D-F) Rescue experiments for the *short-fin* mutant phenotype. (D) *sof* mutant. (E) *sof* mutant expressing wild-type cx43 in keratinocytes. (F) *sof* mutant expressing wild-type cx43 in osteoblasts. Data are presented as mean ± SD (n = 5–10 fish per group). The mean values for WT fish were set to 100. *P < 0.05 versus the corresponding mutant control (*alf* in B and C; *sof* in E and F) (Welch’s one-way ANOVA followed by Dunnett’s multiple-comparisons test). ns, not significant. Black scale bars in whole-body images, 10 mm. White scale bars in fin-ray images, 300 µm.

In the *sof* mutant, expression of wild-type zebrafish cx43 in keratinocytes partially rescued fin size (Fig. 3D, E), suggesting that restoring cx43 function in keratinocytes alone is effective, but insufficient to fully rescue the reduced fin size. By contrast, expression of wild-type cx43 in osteoblasts elongated fin-ray segment beyond the wild-type level without affecting fin size (Fig. 3F). This result indicates that restoring cx43 function in osteoblasts is sufficient to rescue the shortened fin-ray segment phenotype in the *sof* mutant. Together, these rescue experiments indicate that fin size regulation requires bioelectric signaling in multiple cell populations, whereas fin-ray segment length can be restored by bioelectric manipulation in osteoblasts alone in the *alf* and *sof* mutants.

### Bioelectric manipulation in keratinocytes prolongs regenerative fin growth associated with persistent Wnt signaling activity

To examine the effects of keratinocyte-specific bioelectric manipulation on fin size regulation, we observed the regeneration process after caudal fin amputation (Fig. 4A). *Tg(krt4:kcnk5b^W169L^)* transgenic fish regenerated longer caudal fins after amputation. Until 8 days post-amputation (dpa), the transgenic fish showed a similar regeneration rate to that of control fish. After 8 dpa, the regeneration rate of control fish declined, whereas that of the transgenic fish was maintained until 16 dpa, resulting in longer regenerated fins.

**Figure 4.**
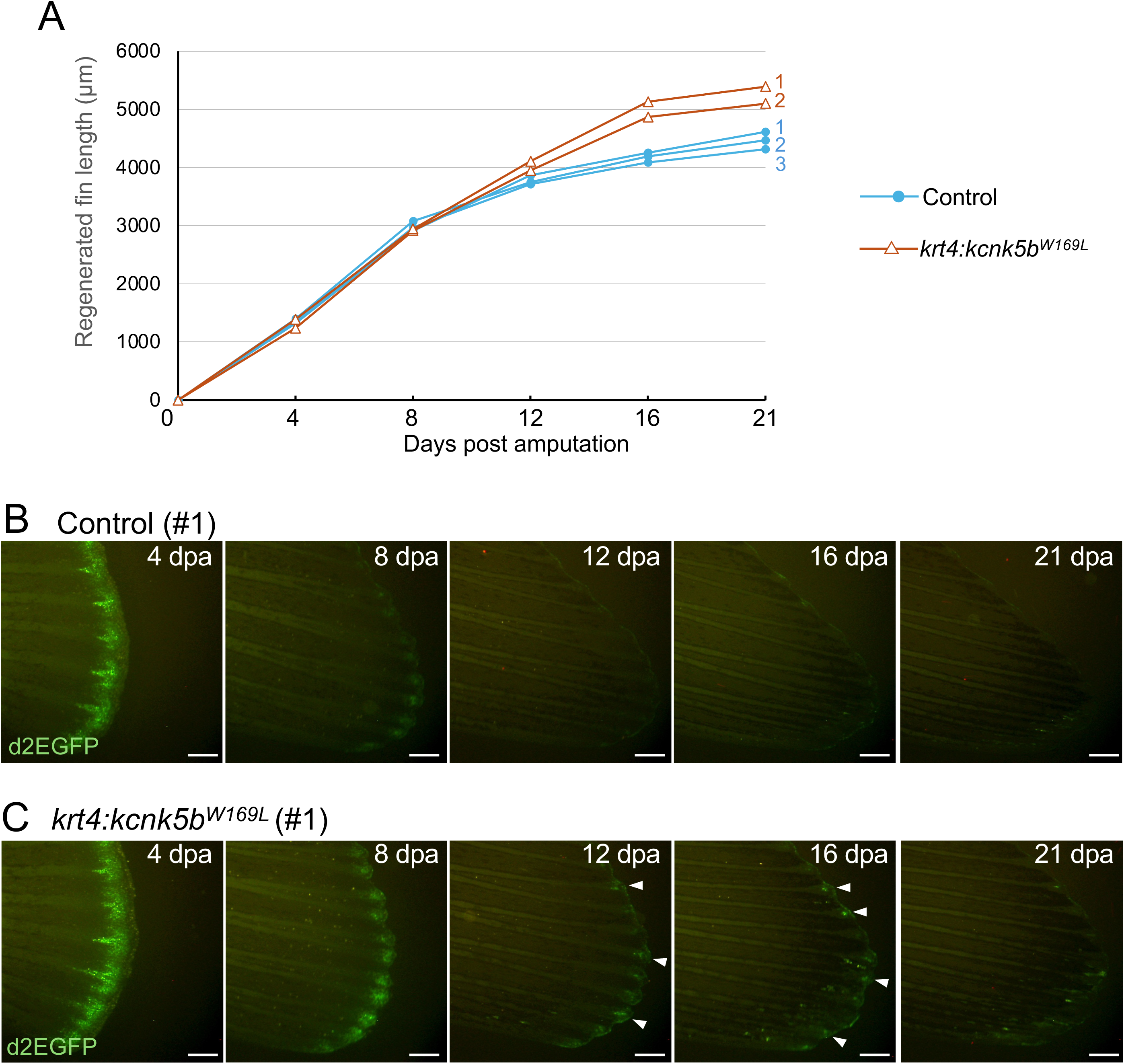
Keratinocyte-specific bioelectric manipulation prolongs fin regeneration with persistent canonical Wnt signaling. (A) Regenerated caudal fin length after amputation. Blue lines and filled circles, control fish. Red lines and triangles, *krt4:kcnk5b^W169L^* transgenic fish expressing kcnk5b^W169L^ in keratinocytes. (B, C) Canonical Wnt signaling activity monitored using the *Tcf/Lef-miniP:dGFP* reporter. (B) Control fish. (C) *krt4:kcnk5b^W169L^* transgenic fish. White scale bar, 300 µm. dpa, days post amputation. d2EGFP, destabilized EGFP.

Because canonical Wnt signaling is known to regulate blastema growth during fin regeneration (Wehner et al., 2014), we monitored canonical Wnt signaling activity during fin regeneration using the *Tcf/Lef-miniP:dGFP* transgenic reporter (Shimizu et al., 2012) (Fig. 4B, C). At 4 dpa, strong reporter activity was observed at the tip of the regenerate in both control and transgenic fish. At 8 dpa, the reporter activity remained at a higher level in the transgenic fish than in the control fish. During 12–16 dpa, the reporter activity was largely absent in the control fish, whereas the reporter activity persisted in the transgenic fish. Notably, sustained regeneration and persistent Wnt signaling reporter activity were observed during the same stage of fin regeneration (8–16 dpa) in the transgenic fish. These findings suggest that keratinocyte-specific bioelectric manipulation is associated with sustained Wnt signaling activity in the blastema during fin regeneration, coinciding with prolonged fin outgrowth.

### Membrane depolarization induces morphological phenotypes opposite to those induced by membrane hyperpolarization

The gain-of-function potassium channel mutant kcnk5b^W169L^ has previously been proposed to hyperpolarize cells expressing it (Perathoner et al., 2014). To test whether direct manipulation of membrane potential can control fin and fin-ray segment morphology, we induced membrane depolarization using the optogenetic tool ChR2, a blue-light-gated cation channel (Zhang et al., 2006). We generated a keratinocyte-specific ChR2-expressing transgenic fish line *Tg(krt4:ChR2^D156C^)* and an osteoblast-specific ChR2-expressing transgenic fish line *Tg(5x osx:ChR2^D156C^)*.

During the optogenetic experiment (Fig. 5A), blue light exposure did not cause morphological changes in the non-transgenic control fish (Fig. 5B, C). In the absence of blue light, the transgenic fish *Tg(krt4:ChR2^D156C^)* and *Tg(5x osx:ChR2^D156C^)* developed normally (Fig. 5D, F). Upon blue-light exposure, ChR2 activation in keratinocytes selectively reduced fin size (Fig. 5E), whereas ChR2 activation in osteoblasts selectively shortened fin-ray segments without affecting overall fin size (Fig. 5G). These results demonstrate that optogenetic membrane depolarization is sufficient to reduce fin size and fin-ray segment length in a cell-type-dependent manner, producing phenotypes opposite to those induced by the kcnk5b^W169L^ transgenic fish.

**Figure 5.**
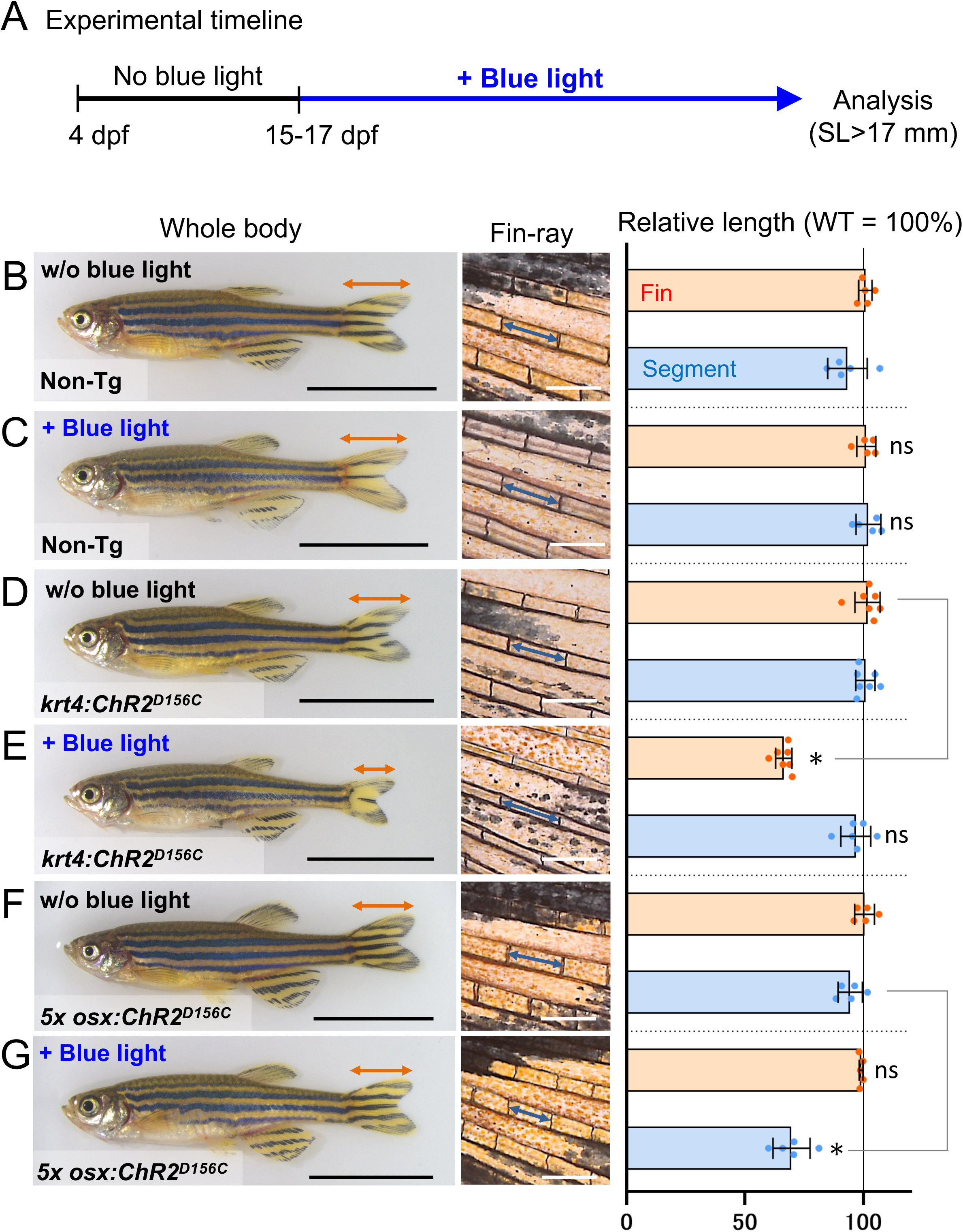
Optogenetic membrane depolarization induced cell-type-specific fin phenotypes. (A) Timeline of the optogenetic experiment. Transgenic fish or non-transgenic control fish were reared under blue-light-blocked conditions from 4 dpf to 15–17 dpf, and subsequently exposed to blue light until the time of analysis. (B, C) Non-transgenic control fish reared without (B) or with blue light (C). (D, E) Transgenic fish expressing ChR2^D156C^ in keratinocytes reared without (D) or with (E) blue light. (F, G) Transgenic fish expressing ChR2^D156C^ in osteoblasts reared without (F) or with (G) blue light. Data are presented as mean ± SD (n = 5–7 fish per group). The mean values for WT fish were set to 100. *P < 0.05 versus the corresponding control without blue light (Student’s t-test or Welch’s t-test). ns, not significant. Black scale bars in whole-body images, 10 mm. White scale bars in fin-ray images, 300 µm. dpf, days post fertilization.

## DISCUSSION

Our study establishes a framework to dissect bioelectric regulation of zebrafish fin morphology and demonstrates that fin size and fin-ray segment length are independently controlled through distinct cellular targets. Bioelectric manipulation in keratinocytes primarily affected fin size, whereas manipulation in osteoblasts selectively altered fin-ray segment length. Combining these manipulations further demonstrated that fin size and fin-ray segment length can be regulated independently despite involving the same bioelectric regulators (Fig. 2).

Previous studies have reported that increased potassium channel activity in fibroblasts or mesenchymal cells induces fin overgrowth (Perathoner et al., 2014; Silic et al., 2020; Stewart et al., 2021). These observations suggested that different bioelectric regulators might act through a common mesenchymal cell lineage. Consistent with these findings, mesenchymal-specific expression of kcnk5b^W169L^ using the *actinodin1^MC^* promoter also reproduced the fin overgrowth phenotype (Table 1, Fig. S2D). However, our study identifies keratinocytes as an additional cellular target of bioelectric fin size regulation. Importantly, rescue experiments further indicate that bioelectric manipulation in keratinocytes alone is insufficient to rescue the fin size phenotype of the *alf* mutant (Fig. 3B), suggesting that multiple cell populations, including mesenchymal cells, contribute to fin size regulation.

Our rescue experiments also showed that restoration of cx43 in keratinocytes alone is insufficient to rescue fin size reduction of the *sof* mutant (Fig. 3E), suggesting that additional cell populations contribute to fin size regulation. Consistent with this finding, a previous study reported that cx43 function in dermal fibroblasts rescues the *sof* mutant phenotype (Dardis et al., 2017). In addition, our osteoblast-specific rescue experiments demonstrate that restoration of cx43 in osteoblasts is sufficient to rescue the fin-ray segment length phenotype of the *sof* mutant (Fig. 3F). Our findings indicate that cx43 function in osteoblast-lineage cells is sufficient to regulate fin-ray segment length.

During fin regeneration, canonical Wnt signaling activity in distal blastemal mesenchymal cells regulates cell proliferation and blastema growth (Wehner et al., 2014). Although a previous study reported that kcnk5b overexpression activates canonical Wnt signaling in a cell-autonomous manner (Yi et al., 2021), our results showed that keratinocyte-specific expression of kcnk5b^W169L^ maintained canonical Wnt signaling activity in the distal blastema, coinciding with sustained blastema growth (Fig. 4B, C). These findings suggest that bioelectric signaling in keratinocytes may indirectly influence Wnt signaling activity in the distal blastema, thereby contributing to prolonged fin outgrowth.

Mutations in different potassium channels (kcnk5b, kcnh2a, and kcnj13) produce similar fin overgrowth phenotypes (Perathoner et al., 2014; Silic et al., 2020; Stewart et al., 2021), suggesting that these phenotypes may depend on potassium permeability rather than the identity of a specific potassium channel. To further test this idea, we expressed human KCNJ2, a potassium channel structurally distinct from kcnk5b, in keratinocytes or osteoblasts (Fig. S7). Cell-type-specific expression of KCNJ2 induced fin size and fin-ray segment length phenotypes similar to those produced by kcnk5b^W169L^. These findings support the idea that increased potassium permeability and the resulting hyperpolarization may contribute to these phenotypes. Our optogenetic experiments further demonstrate that ChR2-mediated membrane depolarization is sufficient to alter fin size and fin-ray segment length (Fig. 5). Membrane depolarization produced morphological phenotypes opposite to those induced by overexpression of potassium channels, supporting the idea that membrane potential functions as a regulatory signal controlling fin morphogenesis in a cell-type-dependent manner.

Our results showed that bioelectric manipulation in epidermal keratinocytes regulates overall fin size. This role is reminiscent of that of the apical ectodermal ridge (AER), a major signaling center controlling limb bud outgrowth during embryogenesis (Fernandez-Teran and Ros, 2008; Yano et al., 2012). A recent study reported that kcnk5b overexpression induced embryonic fin bud overgrowth through membrane depolarization (Jiang et al., 2024). In contrast, our experiments showed that optogenetic membrane depolarization suppressed fin growth during the post-embryonic stage (Fig. 5E). These apparently opposite effects suggest that the developmental responses to bioelectric signaling differ between embryonic limb bud growth and post-embryonic fin growth. Together, these observations suggest that the cellular targets and developmental roles of bioelectric signaling may depend on developmental stage and tissue context.

The independent regulation of appendage size and skeletal element size may have contributed to the morphological diversification of vertebrate appendages during evolution. The limbs of terrestrial vertebrates exhibit remarkable diversity in size and proportion, whereas their fundamental skeletal organization is highly conserved (Mariani and Martin, 2003). In contrast, secondary aquatic vertebrates, including modern cetaceans and several extinct aquatic reptiles, possess enlarged flippers (forelimbs) containing numerous shortened phalanges, a condition known as hyperphalangy (Fedak and Hall, 2004). Because fish paired fin rays and tetrapod digits share a developmental origin (Shimada et al., 2013) and exhibit deep homology in their genetic programs (Nakamura et al., 2016), the independent regulation of fin size and fin-ray segment length observed in zebrafish may provide insight into the diversification of skeletal organization in vertebrate appendages. Consistent with this possibility, our cell-type-specific bioelectric manipulation generated a hyperphalangy-like fin morphology in zebrafish (Fig. 2E, G lane 6). Furthermore, hyperphalangy has been proposed to arise through prolonged appendage outgrowth during limb development (Cooper et al., 2018). Our regeneration experiments suggest that keratinocyte bioelectric signaling can maintain appendage outgrowth (Fig. 4A). Together, these findings provide a concept for considering how independent regulation of appendage growth and skeletal element growth could contribute to evolutionary innovations such as hyperphalangy.

### Study limitations

A limitation of this study is that membrane potential changes induced by the ion-channel manipulations were not directly measured in adult fin tissues. Thus, although the known properties of the channels and the phenotypes induced by ChR2 activation support a role for membrane potential in regulating fin morphology, the magnitude and spatial distribution of membrane potential changes *in vivo* remain to be determined. In addition, although keratinocyte-specific bioelectric manipulation was associated with persistent Wnt signaling activity during fin regeneration, our experiments do not establish a causal relationship between Wnt signaling and the prolonged fin outgrowth.

## MATERIALS AND METHODS

### Zebrafish lines and maintenance

Zebrafish were bred and maintained under standard laboratory conditions (Westerfield, 2007). The Tübingen strain (Tü) was used as the wild-type zebrafish line. The *another-long-fin* mutant was obtained from the Zebrafish International Resource Center (ZIRC). The *short-fin* mutant was kindly provided by Dr M. Kathryn Iovine of Lehigh University. All experiments were approved by the Animal Experiments Committee of Osaka University, Japan.

### Measurement

Before taking images, zebrafish were briefly anaesthetized with MS222 (Sigma-Aldrich, St. Louis, MO, USA). Whole-body images were acquired using an MZ16FA stereomicroscope (Leica, Wetzlar, Germany). Subsequently, fin length and body length (standard length) were measured using ImageJ software (Fig. S1A). The relative fin length was defined as fin length divided by standard length (FL/SL). Fin-ray images and fluorescence images were acquired using a BZ-8000K fluorescent microscope (KEYENCE, Osaka, Japan). Fin-ray segment length was measured using BZ-Analyser software or ImageJ software. Unbranched proximal bone segments of the ventral third (V3) fin-ray were measured, and the mean value for each fish was calculated (Fig. S1B). The resulting values were normalized to the mean value of wild-type fish and presented in each graph.

### Transgenic vector construction

Ubiquitous and cell-type-specific promoters (Dale and Topczewski, 2011; Inohaya et al., 2007; Ju et al., 1999; Knopf et al., 2011; Lalonde et al., 2016; Lee et al., 2014; Mosimann et al., 2011; Pyati et al., 2006; Spoorendonk et al., 2008) were PCR-amplified using KOD plus Neo or KOD Fx Neo DNA polymerase (Toyobo) and cloned into the pTol2 plasmid. To reinforce promoter activity, enhancer-containing regions of the *keratin4*, *actinodin1^MC^*, *RUNX2,* and *osterix* promoters were tandemly aligned (5x promoter). These enhancer-containing regions were predicted based on sequence conservation among fish species using the UCSC Genome Browser (https://genome.ucsc.edu). PCR-amplified enhancer-containing fragments were tandemly inserted upstream region of the original promoter sequence by In-Fusion cloning (Takara Bio). This process is summarized in Figure S4. Zebrafish *kcnk5b* CDS was cloned by RT-PCR, and then the *another-long-fin* mutation, W169L (Perathoner et al., 2014) was introduced by PCR-based site-directed mutagenesis. Similarly, the rat *Cx43* CDS was cloned by RT-PCR, and a dominant-negative mutation, T154A (Beahm et al., 2006) was introduced. The ChR2^D156C^ variant (alias ChR2-XXL) (Dawydow et al., 2014) was generated previously (Aramaki and Kondo, 2020). Sequences of PCR primers are presented in Tables S1-S3.

### Generation of transgenic zebrafish

Transposase mRNA was synthesized *in vitro* and mixed with the Tol2 plasmid DNA. Then, 1 nL of the mixture (10 ng/µL each) was injected into fertilized eggs at the single-cell stage (Kawakami, 2005; Kawakami et al., 2000). Transgene integration was determined by EGFP expression driven by the *ubiquitin-b* promoter, and the expression of mutant genes was confirmed by mCherry or H2B-RFP expression driven by the corresponding cell-type-specific promoter. The phenotypes reported in this study were confirmed in at least two independent stable transgenic lines.

### Optogenetic experiment

The light-gated cation channel ChR2^D156C^ possesses high light responsiveness, high cation permeability, and minimal dependence on exogenous retinal, making it suitable for inducing depolarization in non-neuronal cells. Before blue light exposure, transgenic fish expressing ChR2^D156C^ were reared in red-colored acrylic tanks that block environmental blue light, with circulating water. During blue light stimulation, the transgenic fish were transferred to the stimulation tanks, surrounded by a waterproof LED light strip with 60 blue LED units (JW-System, Tokyo, Japan; product ID: 50B60W66100), and placed in a 28°C incubator. The stimulation tanks were covered with aluminum foil to reflect blue light into the tanks. In the control (non-stimulatory) condition, the transgenic fish were maintained in the red-colored acrylic tanks.

### Statistical analysis

Statistical analyses were performed using GraphPad Prism 8 (GraphPad Software). Data were assessed for normality using the Shapiro–Wilk test. Comparisons among three or more groups were performed using Welch’s one-way analysis of variance (ANOVA), followed by Dunnett’s multiple-comparisons test to compare each experimental group with the corresponding control group. For comparisons between two groups, normally distributed data were analysed using Student’s t-test or Welch’s t-test after assessing equality of variances using the F-test. Differences were considered statistically significant at P < 0.05.

### Immunohistochemistry

Zebrafish were anesthetized briefly for amputation of the caudal fin. Regenerating blastemas were excised at 4 days post amputation and fixed with 4% PFA in PBS overnight at 4 °C. Fixed blastema samples were washed with PBS and incubated in a series of increasing concentrations of sucrose in PBS (5%, 10% and 15% for 1 h at RT, respectively, and 20% overnight at 4 °C). Next, the samples were embedded in O.C.T. Compound (Tissue-Tek) and frozen in liquid nitrogen. Samples embedded in O.C.T. blocks were sectioned at 10 µm thickness using a Leica CM1850 cryostat. The sections were quickly transferred to MAS-coated slides and dried at room temperature. Immunohistochemistry was performed by a conventional method. Anti-DsRed antibody (Takara Bio 632496, 1:200) and Alexa Fluor 594-conjugated anti-Rabbit IgG (invitrogen A11012, 1:500) were used to detect the expression of the reporter H2B-RFP. The section images were acquired using an Axio Observer 7 microscope (Carl Zeiss) and analyzed using ZEN software (Carl Zeiss).

## Declaration of generative AI tools used in the writing process

During the revision of this manuscript, the authors used ChatGPT in order to check grammar and improve readability of the text. After using this tool, the authors reviewed and edited the content as needed and take full responsibility for the content of the publication.

## Acknowledgements

The authors are grateful to Dr M. Kathryn Iovine for providing the *short-fin* mutant and to Dr Matthew P. Harris for comments on early drafts of the manuscript. The authors are also grateful to members of the Kondo laboratory for comments and experimental support; to Yoko Shigeno for experimental assistance; to Dr Seita Miyazawa for the design and production of the red acrylic tank; and to Mana Kadota, Hitomi Tanimoto, and Midori Tanaka for assistance in maintaining the fish facility.

## Competing interests

The authors declare no competing or financial interests.

## Author Contributions

T.A. and S.K. designed research; T.A. generated materials, performed experiments, and analysed data; T.A. and S.K. interpreted results and wrote the manuscript.

## Funding

This work was supported by JSPS KAKENHI JP16K14736, JP20K06655 and JST PREST JPMJPR2042.

## Data and resource availability

All relevant data and details of resources can be found within the article and its supplementary information. Newly generated transgenic zebrafish lines and DNA constructs are available from the corresponding authors upon reasonable request.

## Notes

### Competing Interest Statement

The authors have declared no competing interest.

### Summary of Updates

The manuscript has been substantially revised with additional experiments and analyses to clarify the cell-type-dependent regulation of fin size and fin-ray segment length. Figures and supplementary materials have also been updated.

