## Supplemental Figures and Tables for "Cell-type-dependent bioelectric regulation of fin size and fin-ray segment length in zebrafish"

### SUPPLEMENTARY MATERIALS

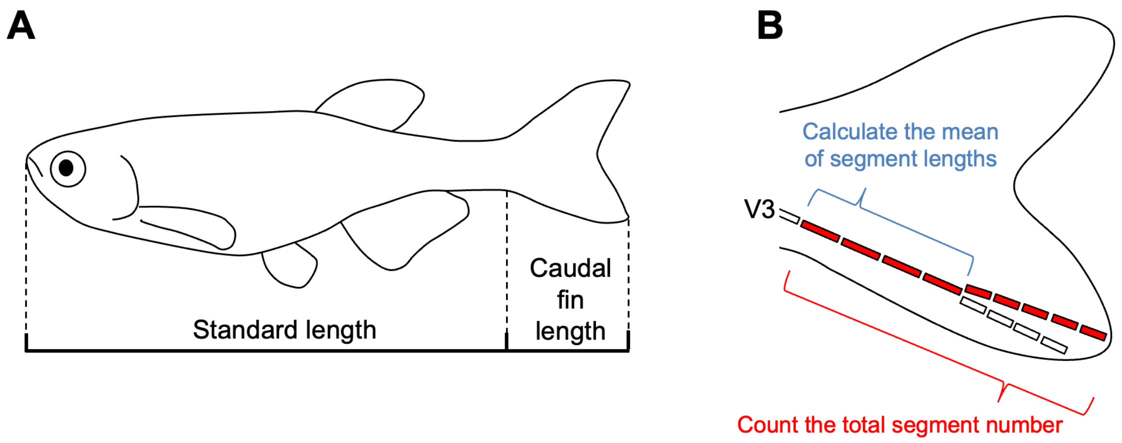


**Figure S1 Measurement of fin length, fin-ray bone segment length, and segment number.**

(A) Measurement method for standard length (SL) and caudal fin length. (B) Measurement method for fin-ray segment length and segment number. Unbranched proximal bone segments of the ventral third (V3) fin-ray were measured, and the mean value was calculated. The total number of segments in the longest V3 fin ray was counted.

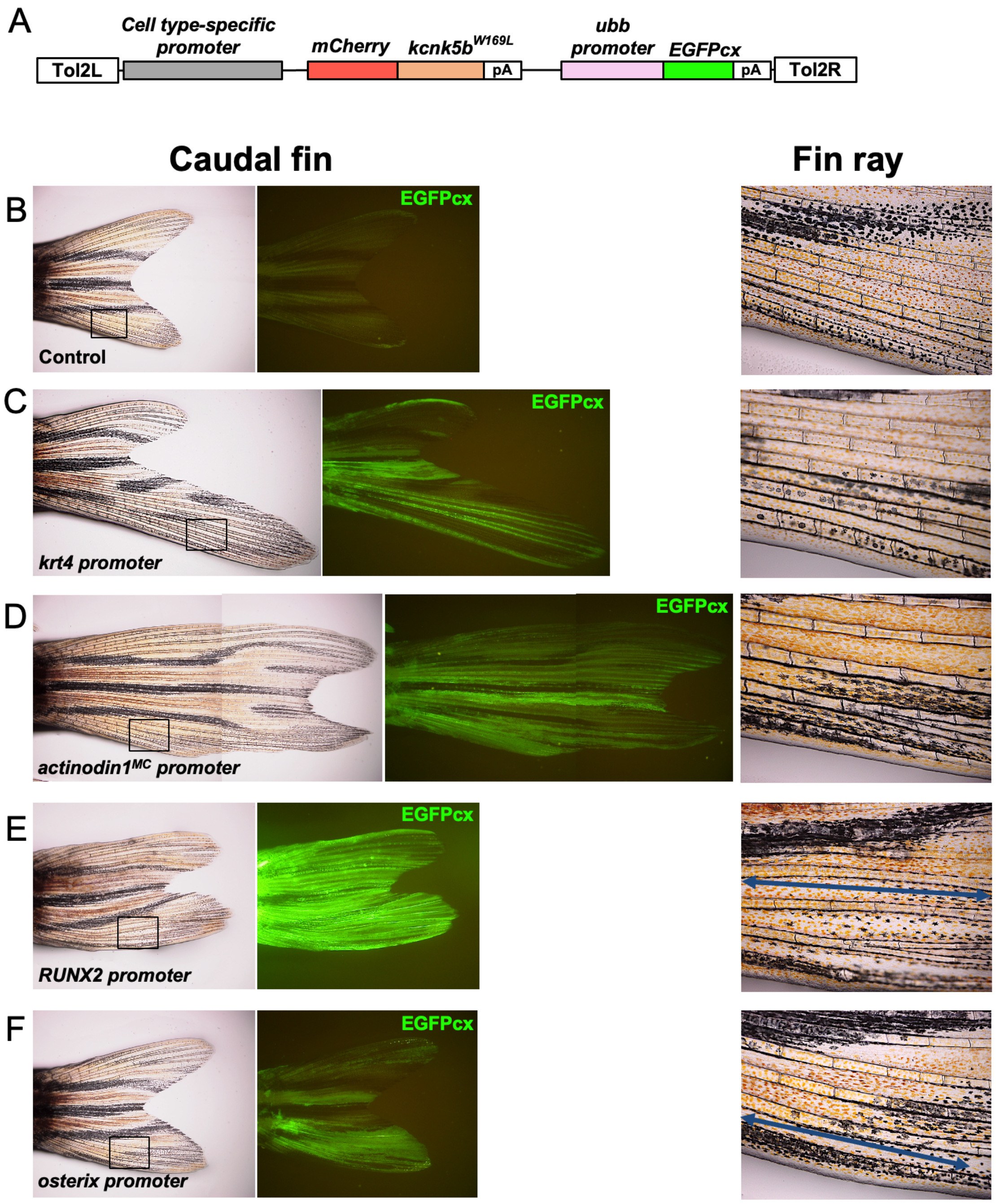


**Figure S2 Typical examples of promoter screening results with kcnk5b$^{W169L}$.**

(A) Schematic diagram of the transgenic construct for the first screening. The promoter region was replaced with various cell-type-specific promoters. (B-F) Caudal fin morphology and fin-ray segments in wild-type fish (B) and F0 transgenic fish expressing kcnk5b$^{W169L}$ under different cell-type-specific promoters (C-F). Transgene integration was confirmed by EGFPcx fluorescence. Areas enclosed by squares in caudal fin images are magnified in the fin-ray images on the right. Blue bars in the fin-ray images indicate elongated fin-ray segments (E, F).

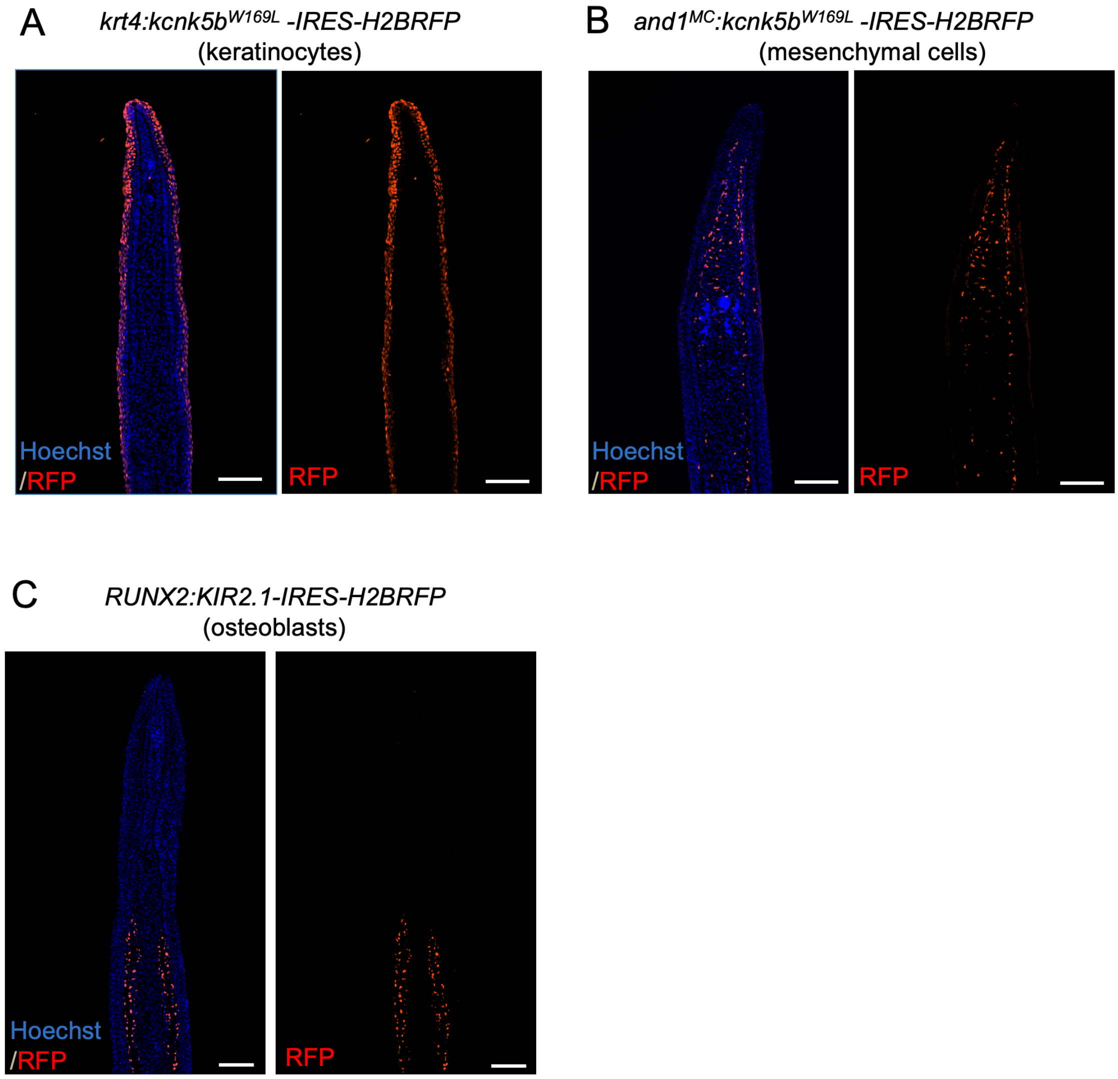


**Figure S3 Confirmation of cell-type-specific promoter activity in the 4-dpa blastema**

(A-C) Immunohistochemical images show specific expression patterns driven by the promoters used in this study. (A) Keratinocyte-specific expression under the *krt4* promoter. (B) Mesenchymal cell-specific expression under the *and1*$^{MC}$ promoter. (C) Osteoblast-specific expression under the *RUNX2* promoter. White scale bars, 100 µm. dpa, days post amputation.

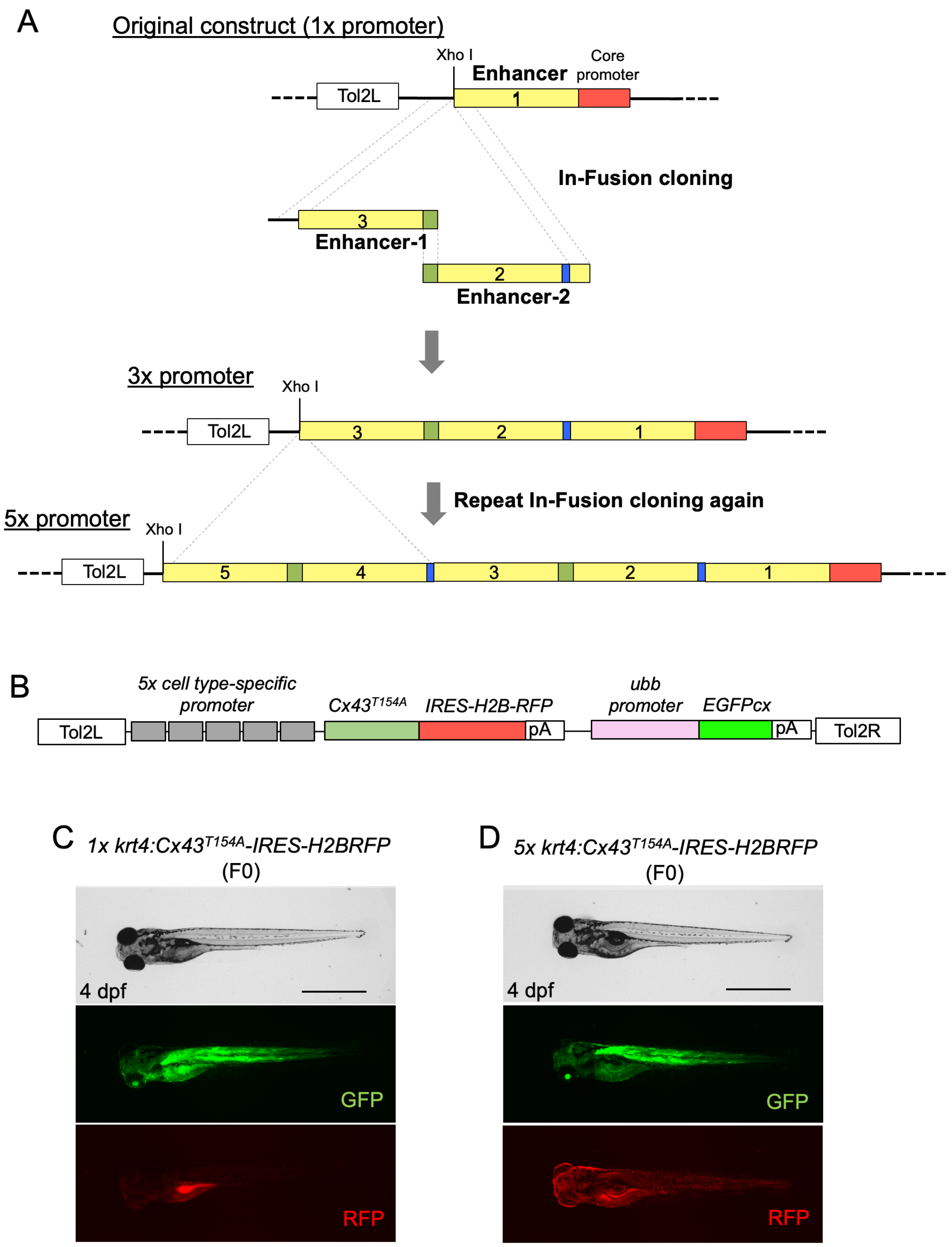


**Figure S4 Generation of multimerized promoter constructs**

(A) Construction process of 5x promoter constructs. Two enhancer-containing DNA fragments (Enhancer-1 and Enhancer-2) were amplified by PCR and inserted upstream of the original (1x) promoter by In-Fusion cloning, resulting in a 3x promoter construct. Then, the same enhancer fragments were inserted again upstream of the 3x promoter, resulting in a 5x promoter construct. (B) Schematic diagram of the transgenic construct for the second screening with $Cx43^{T154A}$. (C, D) Comparison of expression levels between the *1x krt4* promoter (C) and the *5x krt4* promoter (D). Transgene integration was confirmed by GFP fluorescence, and keratinocyte-specific expression was detected by RFP. Black scale bars, 1 mm. dpf, days post fertilization.

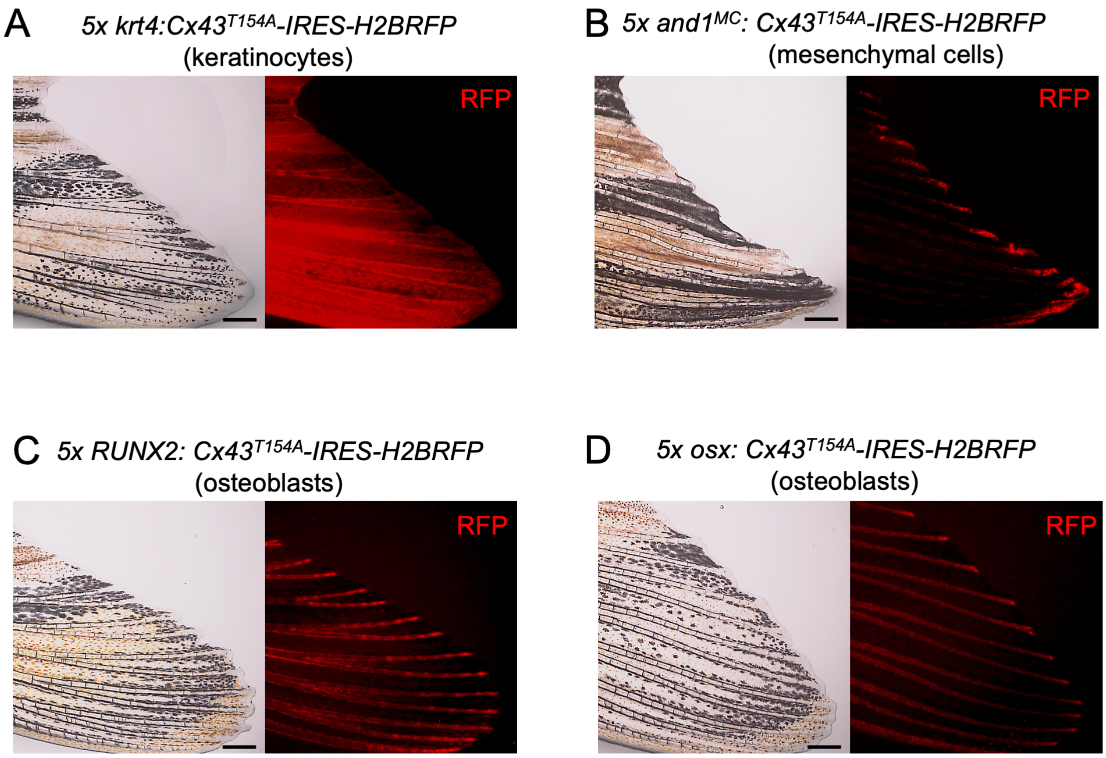


**Figure S5 Confirmation of the cell-type specificity of 5x promoter transgenes**

(A-D) Specific expression under different 5x cell-type-specific promoters in caudal fins. (A) Keratinocyte-specific expression under the *5x krt4* promoter. (B) Mesenchymal cell-specific expression under the *5x and1*$^{MC}$ promoter. (C) Osteoblast-specific expression under the *5x RUNX2* promoter. (D) Osteoblast-specific expression under the *5x osx* promoter. Black scale bars in bright-field images, 300 µm.

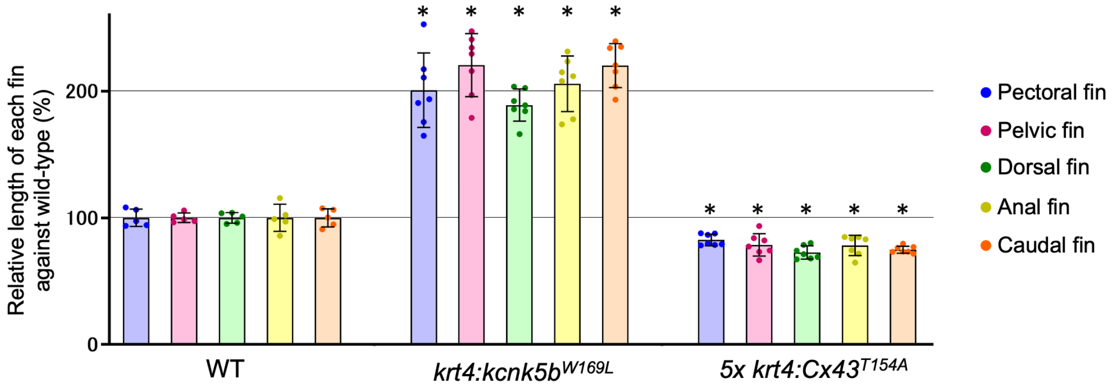


**Figure S6 Effects of keratinocyte-specific bioelectrical manipulations on size in all fins.**

Data are presented as mean ± SD (n = 5–7 fish per group). Mean values for wild-type fish were set to 100 for each fin. *P < 0.05 versus WT (Welch's one-way ANOVA followed by Dunnett's multiple-comparisons test). ns, not significant.

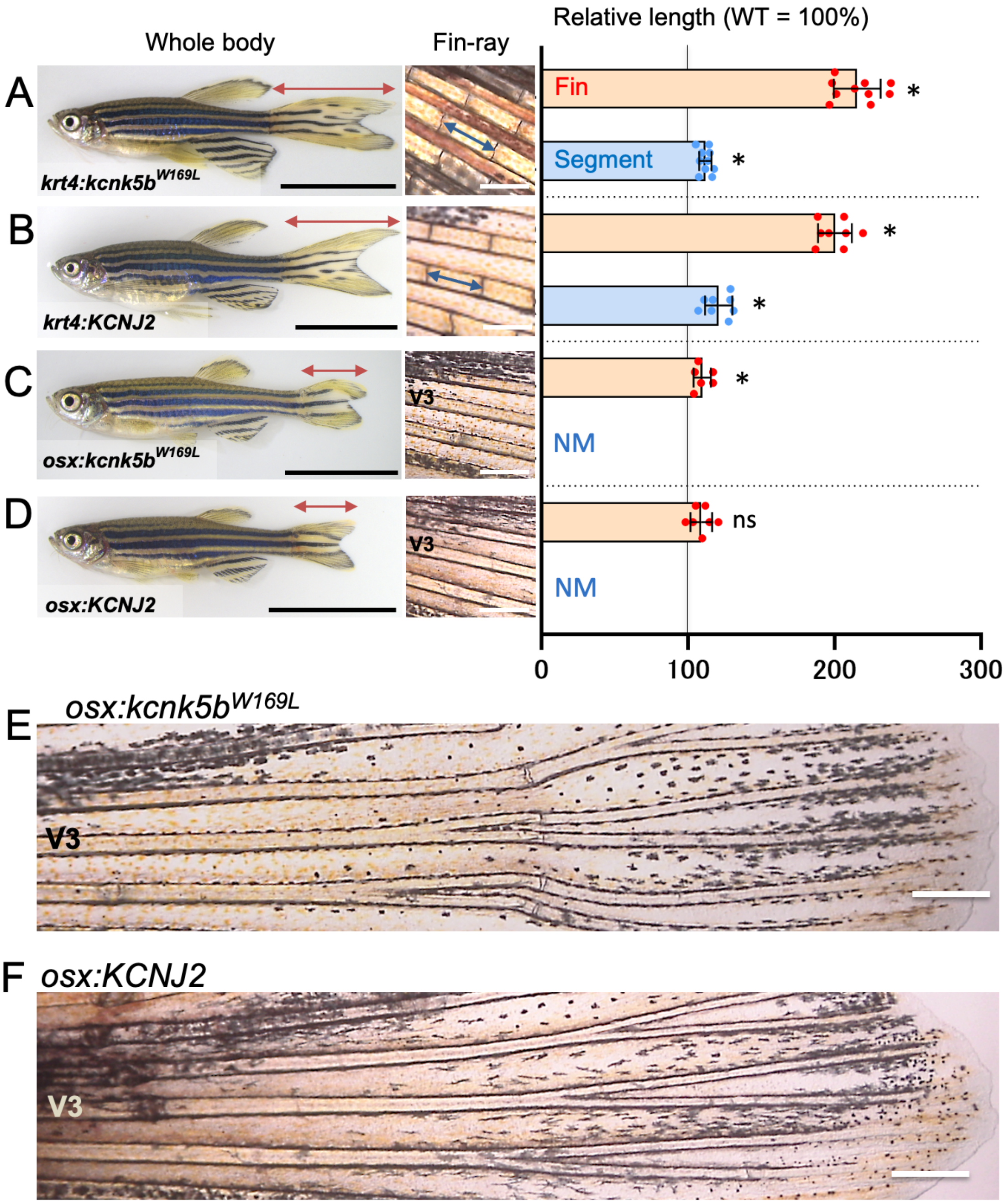


**Figure S7 Expression of different types of potassium channels induces the same fin phenotypes.**

(A) *krt4:kcnk5b*$^{W169L}$ transgenic fish expressing kcnk5b$^{W169L}$ in keratinocytes. (B) *krt4:KCNJ2* transgenic fish expressing KCNJ2 in keratinocytes. (C) *osx:kcnk5b*$^{W169L}$

transgenic fish expressing kcnk5b$^{W169L}$ in osteoblasts. (D) *osx:KCNJ2* transgenic fish expressing KCNJ2 in osteoblasts. (E, F) Images showing a wider field of view of fin-ray images in (C, D). Data are presented as mean ± SD (n = 6–10 fish per group). The mean values for WT fish were set to 100. *P < 0.05 versus WT (Welch's one-way ANOVA followed by Dunnett's multiple-comparisons test). ns, not significant. Black scale bars in whole-body images, 10 mm. White scale bars in fin-ray images, 300 µm. NM, not measurable because fin-ray joints were largely absent, preventing reliable determination of individual segment length.

| Gene | Primer name | Sequence | Reference and remark |
|---|---|---|---|
| ***kcnk5b***$^{W169L}$ | kcnk5b-F_SalI | attgtcgaccaccATGGCAGATAAAGGACCT | Perathoner, 2014 |
| | kcnk5b-R_NotI | attgcggccgcTCAACCAAAGCTTTTCCTC | |
| | kcnk5b W169L-F | CCTTTTATTGGGCTTCTTGGTTCACTT | |
| | kcnk5b W169L-R | AGAAGCCCAATAAAAGGAACACAATTG | |
| ***Cx43***$^{T154A}$ | Rat Cx43-F_SalI | gcaaagaattgtcgaccaccATGGGTGACTGGAGTGCCTT | Beahm, 2006 |
| | Rat Cx43-R_NotI | ttagggggggggggcggccgcTTAAATCTCCAGGTCATCAG | |
| | Rat Cx43 T154A-F | GCTGAGAGCCTACATCATCAGCAT | |
| | Rat Cx43 T154A-R | ATGTAGGCTCTCAGCAAGCCGCCC | |

**Table S1 List of primers used to generate mutant gene constructs.**

Uppercase: sequence for binding to the target, lowercase: linker sequence with restriction enzyme recognition site, underline: mutated sequence.

| Promoter | Primer name | Sequence | Reference and remark |
| --- | --- | --- | --- |
| ***ubiquitin-b*** | ubb pro-F_SpeI | gccactagtACCAGCAAAGTTCTAGAA | Mosimann, 2011 |
| | ubb pro-R_XmaI | attcccgggTCTGTAAACAAATTCAAA | |
| ***keratin1c19e*** | krt1c19e pro-F_NheI | attgctagcTGGCTGAGATGACATGTTTGG | Lee, 2014 |
| | krt1c19e pro-R_AgeI | attaccggtGTGGATGGTGGTTGGTGTCTT | |
| ***keratin4*** | krt4 pro-F_XbaI | atttctagaGCTGCCAAACCCTGTAATGT | Ju, 1999 |
| | krt4 pro-R_AgeI | attaccggtGATGCCTGTGTCTTTGAGTTG | |
| ***col2a1a*** | col2a1a pro-F_NheI | attgctagCCTCTGACACCTGATGCCAATT | Dale, 2011 |
| | col2a1a pro-R_AgeI | attaccggTGCAGGTCCTAAGGGGTGAAAG | |
| ***RUNX2*** **(Human)** | Hunam RUNX2 enh-F_NheI | attgctagcAATTTGGCTCCATGTTTTGGT | Knopf, 2011<br>Human RUNX2 enhancer was linked to minimal promoter (TAGAGGGTATATAATGGAAGCTCG ACTTCCAG). |
| | Human RUNX2 enh-R_SmaI | attcccgGGCAGGCAGTAGATGTGTGAG | |
| ***osterix*** **(Medaka)** | Medaka osx pro-F_NheI | attgctagcTGAACATGTCAGTGCCATCA | Spoorendonk, 2008 |
| | Medaka osx pro-R_AgeI | attaccggtCGGGACAGTTTGGAAGAAGT | |
| ***osteocalcin*** **(Medaka)** | Medaka ostcn pro-F_NheI | attgctagcATTAGTGGTGCAGCGCTGCCA | Inohaya, 2007 |
| | Medaka ostcn pro-R_AgeI | attaccggtGATGCTTCGCAGTCCGTTAAT | |
| ***actinodin1***$^{MC}$ | and1 pro-F_NheI | cgagctagcTTACAGCTTTAAGACACCTCTA | Lalonde, 2016<br><br>Mesenchymal cell specific promoter: Epi Fragment was deleted by PCR. |
| | and1 pro-R_AgeI | attaccggtGATTCCTTTGGATGAAATTAATTAC | |
| | and1 (Δepi)-F | ACATTTTTTTAACATAAAGCACAGATGTTT | |
| | and1 (Δepi)-R | TATGTTAAAAAAATGTGGAAACATCTGG | |
| ***msxb*** | msxb pro-F_NheI | attgctagcTGTGTAGTATAAAACCAGGT | Pyati, 2006 |
| | msxb pro-R_SalI | attgtcgacGGCGTTGGGCTCGGGTGAC | |
| ***acp5a*** | acp5a pro-F_NheI | attgctagcATGTAACTTCCAACACTCAAT | This study |
| | acp5a pro-R_AgeI | attaccggtCTACAAAACAACATACAAACA | |

**Table S2 List of primers used to clone cell-type-specific promoters.**

Uppercase: sequence for binding to the target, lowercase: linker sequence with restriction enzyme recognition site.

| 5x promoter | Fragment | Primer name | Sequence |
|---|---|---|---|
| 5x *keratin4* promoter | 1x krt4 promoter | Same as shown in supplementary table S2. | |
| | krt4 enhancer-1 | Tol2L-F | ACTTGGGCCCGGCTCGA |
| | | krt4 enh-R_U15 | cagtaagatcaaaggCACACTCTTGAGGGGGTG |
| | krt4 enhancer-2 | krt4 enh-F_U15 | cctttgatcttactgGCTGCCAAACCCTGTAAT |
| | | krt4 enh-R_krt4 | gtttggcagctctagcACACTCTTGAGGGGGTG |
| 5x *RUNX2* promoter | 1x RUNX2 promoter | Same as shown in supplementary table S2. | |
| | RUNX2 enhancer-1 | Tol2L-F | ACTTGGGCCCGGCTCGA |
| | | RUNX2 enh-R_U15 | cagtaagatcaaaggGGCAGGCAGTAGATGT |
| | RUNX2 enhancer-2 | RUNX2 enh-F_U15 | cctttgatcttactgAATTTGGCTCCATGTTTT |
| | | RUNX2 enh-R_RUNX2 | catggagccaaattgGGCAGGCAGTAGATGT |
| 5x *osterix* promoter | 1x osterix promoter (3 kbp) | Medaka osx pro (3k)-F_NheI | cccggctcgagctagcATTGAAGTGCGCAGTTG |
| | | Medaka osx pro-R_AgeI | attaccggtCGGGACAGTTTGGAAGAAGT |
| | osx enhancer-1 | Tol2L-F | ACTTGGGCCCGGCTCGA |
| | | osx enh-R_U15 | cagtaagatcaaaggATAAAAAAGAATCAGGGATGGA |
| | osx enhancer-2 | osx enh-F_U15 | cctttgatcttactgATTGAAGTGCGCAGTTG |
| | | osx enh-R_osx | ctgcgcacttcaatgATAAAAAAGAATCAGGGATGGA |

**Table S3 List of primers used to generate multimerized promoter constructs.**

Uppercase: sequence for binding to the target, lowercase: linker sequence, underline: homology sequence for In-Fusion cloning.
